# CRISPR/Cas9-Mediated Knockout of ZFP36L1 Impairs Cell Proliferation, Alters Cell-Cycle Progression, and Enhances DNA Damage Responses in MDA-MB-231 Triple-Negative Breast Cancer Cells

**DOI:** 10.64898/2026.08.20.746080

**Authors:** Hope H.G Gandu, Purity T. Y Gandu, Efe Okorare, Michael Uzorchukwu Ochem, Nneamaka Henrietta Okeke, Divine O. Nwachi, Dennis Kure Yusuf, Nwanneka Grace Anene, Reem G A Hamed, Usman Karuma Shuaib

**Affiliations:** University of Westminster, London. United Kingdom; Tambov state university, Tambov. Russia; Gloucestershire Royal Hospital, Gloucestershire. England; Margaret Lawrence University, Abuja. Nigeria; Glangwilli General hospital, Wales. United Kingdom; Benjamin S Carson School of Medicine, Babcock University, Ogun. Nigeria; Abubakar Tafawa Balewa University, Bauchi. Nigeria; University of Nigeria Teaching Hospital, Ituku-Ozalla, Enugu. Nigeria; Alzaiem Alazhari University, Khartoum Bahri. Sudan; Department of Medical Biochemistry, University of Abuja. Nigeria

**Keywords:** CRISPR/Cas9, DNA damage response, Doxorubicin, Genomic stability, MDA-MB-231, Triple-negative breast cancer, ZFP36L1, γ-H2AX

## Abstract

**Background:** Zinc finger protein 36-like 1 (ZFP36L1) is an AU-rich element-binding RNA-binding protein that regulates post-transcriptional gene expression and has been implicated in tumor progression, cell-cycle regulation, and DNA damage responses. However, its functional role in triple-negative breast cancer (TNBC) remains poorly understood. This study investigated the effects of CRISPR/Cas9-mediated ZFP36L1 knockout on cell proliferation, doxorubicin (DOX) sensitivity, cell-cycle progression, and DNA damage responses in MDA-MB-231 TNBC cells.

**Methods:** Wild-type (WT) and CRISPR/Cas9-generated ZFP36L1 knockout (KO) MDA-MB-231 cells were cultured under standard conditions. Cellular proliferation was evaluated by cell counting over three weeks. Cell viability following DOX treatment was determined using the MTT assay, and half-maximal inhibitory concentration (IC_50_) values were calculated. Cell-cycle distribution was assessed by propidium iodide flow cytometry after 24 h of DOX exposure, while DNA damage was quantified by γ-H2AX flow cytometric analysis. Statistical significance was determined using Student’s *t*-test with *P* < 0.05 considered significant.

**Results:** ZFP36L1 knockout reduced the proliferative capacity of MDA-MB-231 cells compared with WT cells. Both cell lines exhibited dose-dependent decreases in viability following DOX treatment. KO cells demonstrated a higher mean IC_50_ than WT cells (9.64 vs. 8.40 μM), indicating a trend toward reduced DOX sensitivity; however, this difference was not statistically significant (*P* = 0.569). Flow cytometric analysis revealed enhanced accumulation of KO cells in the S and G_2_/M phases following DOX treatment, suggesting altered cell-cycle checkpoint regulation. Furthermore, KO cells exhibited elevated basal γ-H2AX expression and greater DOX-induced γ-H2AX accumulation than WT cells, indicating increased DNA damage and impaired maintenance of genomic stability.

**Conclusions:** CRISPR/Cas9-mediated loss of ZFP36L1 suppresses proliferation, alters cell-cycle checkpoint dynamics, and enhances DNA damage accumulation in MDA-MB-231 TNBC cells. These findings indicate that ZFP36L1 plays a context-dependent role in regulating genomic stability and cellular responses to genotoxic stress, highlighting its potential as a biomarker and therapeutic target in triple-negative breast cancer.

## 1. INTRODUCTION

Breast cancer remains the most commonly diagnosed malignancy among women globally, with triple-negative breast cancer (TNBC) accounting for approximately 15–20% of cases and conferring the poorest prognosis due to the absence of estrogen receptor, progesterone receptor, and human epidermal growth factor receptor 2 expression (Alkaraki et al., 2020). The MDA-MB-231 cell line, established from a TNBC patient with an aggressive basal-like phenotype, serves as a principal preclinical model for investigating breast cancer biology, therapeutic resistance, and DNA damage responses (Alkaraki et al., 2020). Given the limited targeted therapeutic options for TNBC, elucidating the post-transcriptional regulatory networks that govern proliferation and chemotherapy sensitivity is essential for identifying novel biomarkers and therapeutic targets.

At the post-transcriptional level, RNA-binding proteins (RBPs) of the tristetraprolin (TTP) family, including zinc finger protein 36 like 1 (ZFP36L1), have emerged as critical regulators of mRNA stability, cellular stress responses, and cancer progression (Saini et al., 2020). ZFP36L1 contains highly conserved tandem CCCH zinc-finger motifs that recognize adenylate-uridylate–rich elements (AREs) within the 3′-untranslated regions (3′UTRs) of target mRNAs, recruiting RNA degradation complexes to mediate transcript decay (Loh et al., 2020). Through this mechanism, ZFP36L1 functions as a post-transcriptional gatekeeper of oncogenic signaling. In silico analyses of primary tumor cohorts have revealed that ZFP36L1 is frequently mutated, epigenetically silenced via enhancer hypermethylation, and significantly downregulated in bladder, breast, and liver cancers (Loh et al., 2020; Chen et al., 2022). Functionally, forced expression of ZFP36L1 in cancer cells markedly suppresses proliferation and tumor growth in vitro and in vivo by destabilizing key oncogenic transcripts involved in hypoxia signaling (HIF1A), cell cycle progression (CCND1, E2F1), and DNA repair (DCLRE1C), whereas its silencing enhances tumor cell growth (Loh et al., 2020; Zhuang et al., 2025). Conversely, combined germline deletion of ZFP36L1 and its paralog ZFP36L2 in murine lymphocytes drives T-cell acute lymphoblastic leukemia via Notch1 stabilization, underscoring the essential role of ZFP36L1 in maintaining hematopoietic genomic homeostasis (Saini et al., 2020).

Nevertheless, the functional classification of ZFP36L1 as a canonical tumor suppressor has been challenged by recent context-specific findings. In chronic myeloid leukemia, CRISPR-Cas9-mediated ZFP36L1 knockout paradoxically decreased cell expansion and upregulated the cyclin-dependent kinase inhibitor CDKN1A via direct 3′UTR binding, indicating that ZFP36L1 can negatively regulate tumor suppressor transcripts in a cell-type-dependent manner (Kaehler et al., 2021). Similarly, in muscle-invasive bladder cancer, ZFP36L1 suppresses colony formation and self-renewal while simultaneously promoting epithelial-mesenchymal transition and invasion through metastasis-associated pathways (Yuan et al., 2022). These conflicting observations highlight that the oncogenic or tumor-suppressive function of ZFP36L1 is contingent upon the cellular transcriptome and signaling environment, necessitating cancer type-specific functional dissection.

The clustered regularly interspaced short palindromic repeats (CRISPR)-CRISPR-associated protein 9 (Cas9) system has revolutionized functional cancer genomics by enabling precise, efficient gene knockout to dissect tumor suppressor and oncogene function with minimal off-target effects when appropriately validated (Chehelgerdi et al., 2024; Karn et al., 2022). In breast cancer research, CRISPR-Cas9-mediated gene editing has been employed to target chemoresistance mechanisms, DNA repair pathways, and cell cycle regulators, providing a robust platform for preclinical therapeutic discovery (Rehman & Abbas, 2026). Doxorubicin (DOX), an anthracycline topoisomerase II inhibitor, remains a cornerstone of neoadjuvant and adjuvant TNBC chemotherapy. Its cytotoxicity is mediated through DNA double-strand break (DSB) induction, reactive oxygen species generation, and activation of p53-dependent cell cycle arrest and apoptosis (Alkaraki et al., 2020). Standard modalities for quantifying DOX-induced genotoxic stress include cell viability assays, cell cycle distribution analysis by propidium iodide (PI) staining, and flow cytometric detection of phosphorylated histone H2AX (γH2AX), a well-established surrogate marker of DSBs and replication stress.

Despite accumulating evidence implicating ZFP36L1 as a post-transcriptional regulator of oncogenic signaling, its specific contributions to TNBC proliferation, doxorubicin sensitivity, and DNA damage checkpoint engagement remain incompletely characterized. The present study employed CRISPR-Cas9-mediated ZFP36L1 knockout in MDA-MB-231 cells to investigate the effects of ZFP36L1 loss on baseline cell proliferation, DOX-induced cytotoxicity, cell cycle distribution, and DNA damage accumulation. We hypothesized that ZFP36L1 disruption would alter proliferative capacity and modulate the cellular response to DOX-induced genotoxic stress, thereby informing the context-dependent role of this RBP in TNBC biology.

## 2. METHODOLOGY

### 2.1 Cell Culture

Human triple-negative breast cancer cell lines MDA-MB-231 wild-type (WT) and CRISPR-Cas9 ZFP36L1 knockout (KO) cells were maintained in Dulbecco’s Modified Eagle Medium (DMEM) supplemented with 10% fetal calf serum (FCS) and 1% penicillin/streptomycin (complete medium) at 37°C in a humidified 5% CO₂ incubator. The ZFP36L1 KO cells were generated previously in the laboratory using CRISPR-Cas9 gene editing and single-cell cloning. Routine subculture was performed upon reaching 85–90% confluency. Briefly, culture medium was aspirated, cell monolayers were rinsed with 5 mL phosphate-buffered saline (PBS), and 2.5 mL trypsin-EDTA was added. After incubation at 37°C for up to 5 min, detached cells were collected, transferred to sterile centrifuge tubes, and washed by centrifugation at 1,500 rpm for 5 min at 25°C. Cell pellets were resuspended in fresh complete medium, and viable cells were enumerated using a hemocytometer. Cell passage numbers were recorded for all experiments. Where indicated, cells were treated with doxorubicin (DOX) to induce replication stress.

### 2.2 Cell Counting and Seeding

Cell suspensions were gently pipetted to ensure homogeneity. A 10 µL aliquot was loaded onto a clean hemocytometer with a coverslip, and cells within the four large corner squares were counted under an inverted microscope (10× or 20× objective) using a consistent counting rule: cells touching the top and left borders were included, whereas those touching the bottom and right borders were excluded. The cell concentration was calculated as:

Cells/mL = (Average count per square)×10,000×Dilution factor

For routine maintenance and proliferation tracking, cells were split at a 1:4 ratio into T25 flasks containing 5 mL complete medium, seeded at approximately 1×10^5^ cells/mL, and incubated under standard conditions. Population doublings were monitored over a three-week period.

### 2.3 MTT Cell Viability Assay

Cell metabolic activity and viability were assessed using the 3-(4,5-dimethylthiazol-2-yl)-2,5-diphenyltetrazolium bromide (MTT) assay (Meerloo et al., 2011). Wild-type and ZFP36L1 KO MDA-MB-231 cells were seeded at 6,000 cells per well in 100 µL complete medium in 96-well plates and allowed to attach overnight. Cells were then treated with serial dilutions of DOX (0.14, 0.28, 0.56, 1.13, 2.25, 4.5, 9.0, and 18.0 µM) for 48 h. Following treatment, the medium was removed, and 100 µL of 0.5 mg/mL MTT reagent in serum-free DMEM was added to each well. After 3–4 h incubation at 37°C, the MTT solution was aspirated, and the resulting formazan crystals were solubilized in dimethyl sulfoxide (DMSO). Plates were agitated on a shaker for 5 min, and absorbance was measured spectrophotometrically at 570 nm using a microplate reader. The half-maximal inhibitory concentration (IC_50_) was determined by linear regression analysis using the equation Y=mx+c , where Y=50 , m is the slope, x represents the IC_50_, and c is the y-intercept. Experiments were conducted in biological quadruplicate (n=4 ).

### 2.4 Cell Cycle Analysis

Cell cycle distribution was analyzed by PI staining and flow cytometry. Cells were seeded in T25 flasks and treated with DOX at 0.005 µM or 0.5 µM for 24 h. Cells were harvested, washed twice with ice-cold PBS, and fixed overnight in ice-cold 70% ethanol at 4°C. Fixed cells were washed with PBS, stained with PI (1 mg/mL) in the presence of RNase A (1%) for 30 min at room temperature in the dark, and analyzed by flow cytometry. Histograms of DNA content were deconvoluted to determine the percentage of cells in G0/G1, S, and G2/M phases.

### 2.5 DNA Damage Assay: γH2AX Flow Cytometry

DNA double-strand breaks were quantified by flow cytometric detection of phosphorylated histone H2AX (γH2AX). Wild-type and ZFP36L1 KO cells were treated with or without 0.5 µM DOX for 24 h. Cells were collected, washed with PBS, fixed with BD Cytofix Fixation Buffer (cat. no. 554655), and permeabilized with BD Phospho Perm Buffer (cat. no. 558050). Cells were incubated for 30 min with 5 µL anti-γH2AX antibody and 5 µL isotype control antibody. Samples were washed twice with 5 mL BD Stain Buffer, centrifuged at 1,500 rpm for 5 min at 4°C, and resuspended in 0.5 mL Stain Buffer. γH2AX fluorescence was detected in the PE channel using a flow cytometer, and mean fluorescence intensity (MFI) values were recorded for each condition.

### 2.6 Statistical Analysis

Data were analyzed using Microsoft Excel. Differences in cell viability and IC_50_ values between wild-type and KO cells were compared using Student’s t -test; a paired t -test was applied where appropriate for matched experimental conditions. Statistical significance was set at p<0.05.

## 3. RESULT

### 3.1 Loss of ZFP36L1 reduces the proliferative capacity of MDA-MB-231 cells

Cell proliferation was assessed by comparing the growth of wild-type (WT) and ZFP36L1 knockout (KO) MDA-MB-231 cells over a 96-hour culture period. WT cells consistently exhibited higher cell counts than KO cells across all observation periods (Table 1), indicating that disruption of ZFP36L1 expression was associated with reduced cellular proliferation. WT cell counts increased from **1.0 × 10⁶ cells/mL** during the first week to **1.6 × 10⁶ cells/mL** by the third week, whereas KO cells increased from **5.0 × 10⁵ cells/mL** to **1.2 × 10⁶ cells/mL** over the same period. These findings demonstrate a sustained reduction in the growth rate of ZFP36L1-deficient cells compared with WT cells.

**Table 1.** Growth characteristics of wild-type and ZFP36L1 knockout MDA-MB-231 cells after 96 h of culture.

| Weeks | WT cells/ml | KO cells/ml |
| --- | --- | --- |
| 1 | 1000,000 | 500,000 |
| 2 | 870,000 | 630,000 |
| 3 | 1,600,000 | 1200,000 |

WT: wild-type, KO: knockout

Values represent cell counts measured after 96 h of culture. WT = wild-type; KO = ZFP36L1 knockout.

### 3.2 ZFP36L1 knockout cells exhibit reduced sensitivity to doxorubicin

Cell viability following exposure to increasing concentrations of doxorubicin (DOX) was evaluated using the MTT assay. Both WT and KO cells exhibited a dose-dependent reduction in viability with increasing DOX concentration (Figure 1). However, WT cells demonstrated a greater decline in viability than KO cells across most concentrations tested, indicating a trend toward increased sensitivity to DOX in WT cells.

**Figure 1.**
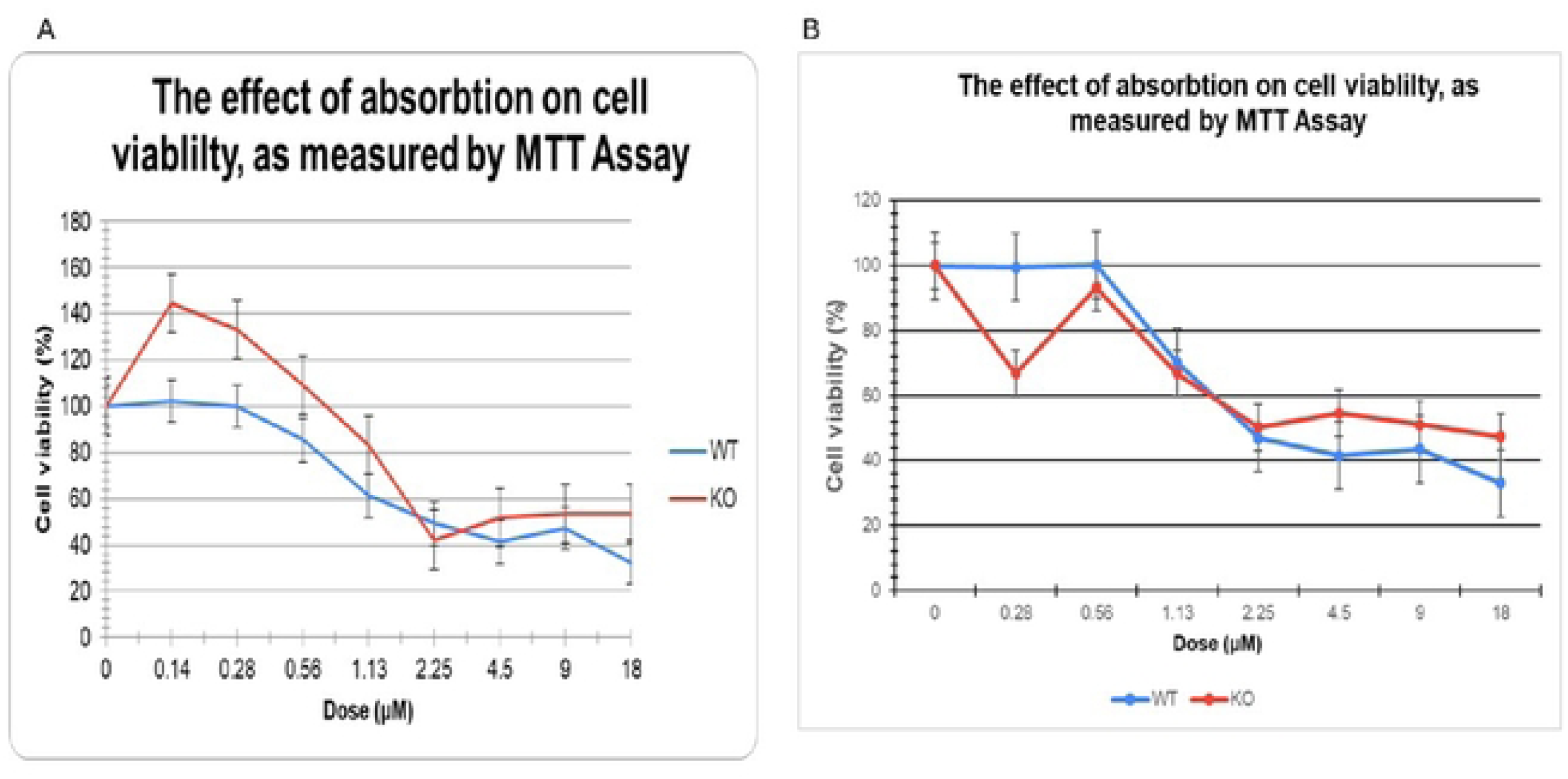
Dose-response curves showing the effect of doxorubicin on the viability of WT and ZFP36L1 knockout MDA-MB-231 cells. Panel (A) shows the 1^st^ experiment and (B) shows the 2^nd^ experiment.

The calculated IC_50_ values further supported this observation. The mean IC_50_ was **8.40 μM** for WT cells and **9.64 μM** for KO cells (Table 2), indicating that a higher concentration of DOX was required to reduce the viability of KO cells by 50%. However, comparison of IC_50_ values between WT and KO cells did not reveal a statistically significant difference (**P = 0.569**).

**Table 2.** IC_50_ values of doxorubicin in WT and ZFP36L1 knockout MDA-MB-231 cells.

| WT cells | KO cells |
| --- | --- |
| **13.00468456** | 8.476668405 |
| **7.462104423** | 10.2935591 |
| **7.725104353** | 10.3161677 |
| **5.39520161** | 9.474864939 |

| Average IC50 value |  |
| --- | --- |
| 8.396773737 | 9.640315036 |

### 3.3 Loss of ZFP36L1 alters cell-cycle progression following doxorubicin treatment

To investigate the effect of ZFP36L1 depletion on cell-cycle progression, WT and ZFP36L1 knockout (KO) MDA-MB-231 cells were analyzed by flow cytometry following 24 h exposure to doxorubicin (DOX). Cell-cycle analysis demonstrated that DOX treatment altered the distribution of cells across the G₀/G₁, S, and G_2_/M phases in both WT and KO cell populations (Figure 2). Compared with untreated controls, DOX-treated cells exhibited a reduction in the proportion of cells in the G₀/G₁ phase, accompanied by an accumulation of cells in the S and G_2_/M phases.

**Figure 2.**
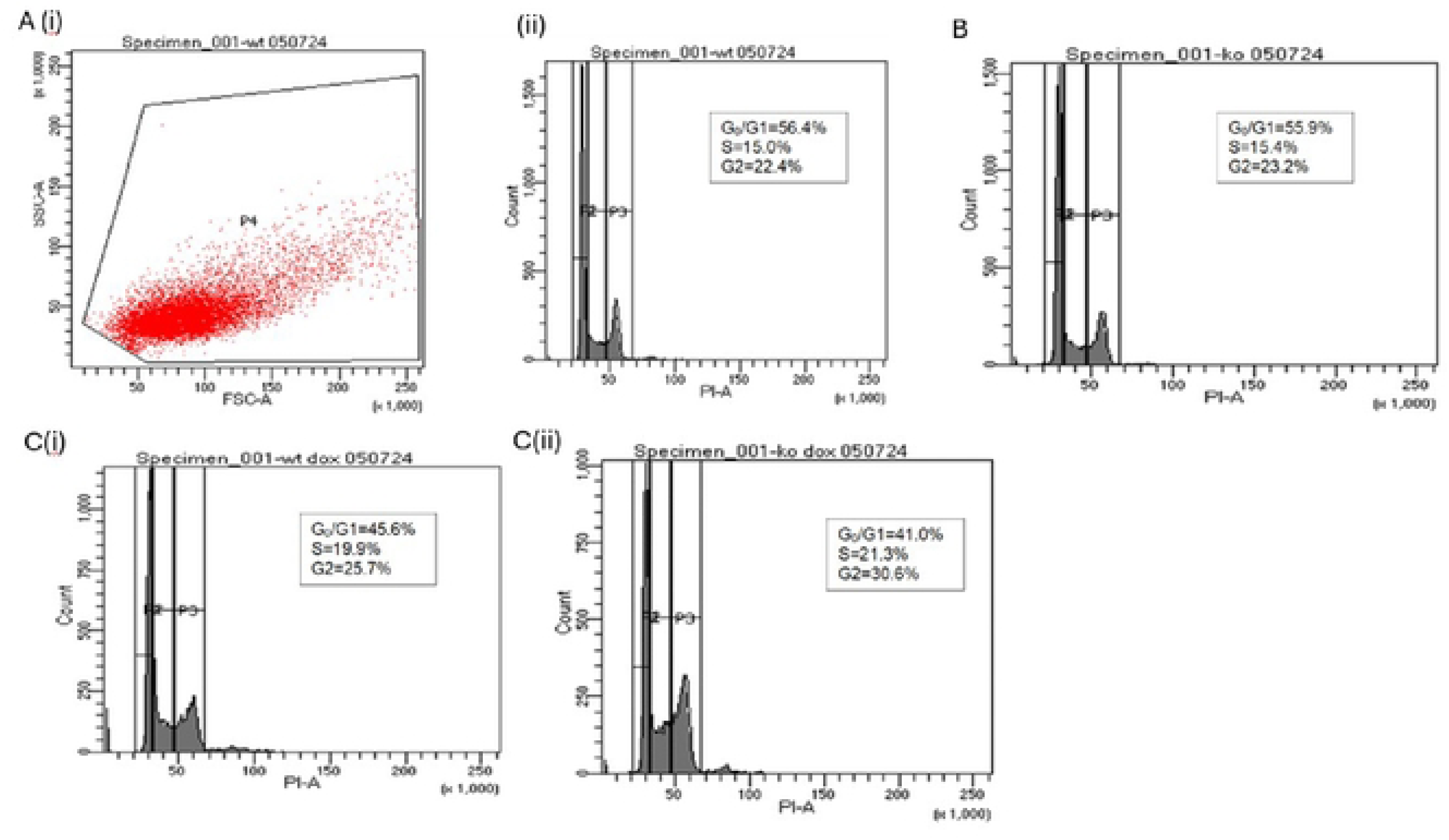
Effect of 0.5 μM doxorubicin on cell-cycle distribution in wild-type and ZFP36L1 knockout MDA-MB-231 cells. Representative flow cytometry histograms showing the distribution of WT and ZFP36L1 knockout (KO) MDA-MB-231 cells across the G₀/G₁, S, and G_2_/M phases following treatment with 0.5 μM doxorubicin for 24 h. Cells were fixed in 70% ethanol, stained with propidium iodide (PI) in the presence of RNase A, and analyzed by flow cytometry. DNA content is presented on the x-axis and cell count on the y-axis. Panel A(i) shows side scatter and forward scatter of granularity; (ii) shows WT cells before DOX treatment; B shows cells before DOX treatment; C(i) shows WT cells after DOX treatment and C(ii) shows KO cells after DOX treatment.

Notably, KO cells displayed a greater proportion of cells arrested in the S and G_2_/M phases than WT cells following treatment with 0.5 μM DOX, suggesting enhanced disruption of cell-cycle progression in the absence of ZFP36L1 (Figure 2). This observation was further supported in a repeat experiment using both 0.5 μM and 0.005 μM DOX (Figure 3). At the lower concentration (0.005 μM), KO cells demonstrated a more pronounced accumulation in the S and G_2_/M phases compared with WT cells, whereas similar trends were observed between the two cell lines at 0.5 μM DOX. Collectively, these findings indicate that loss of ZFP36L1 influences cell-cycle dynamics in response to DOX-induced replication stress.

**Figure 3.**
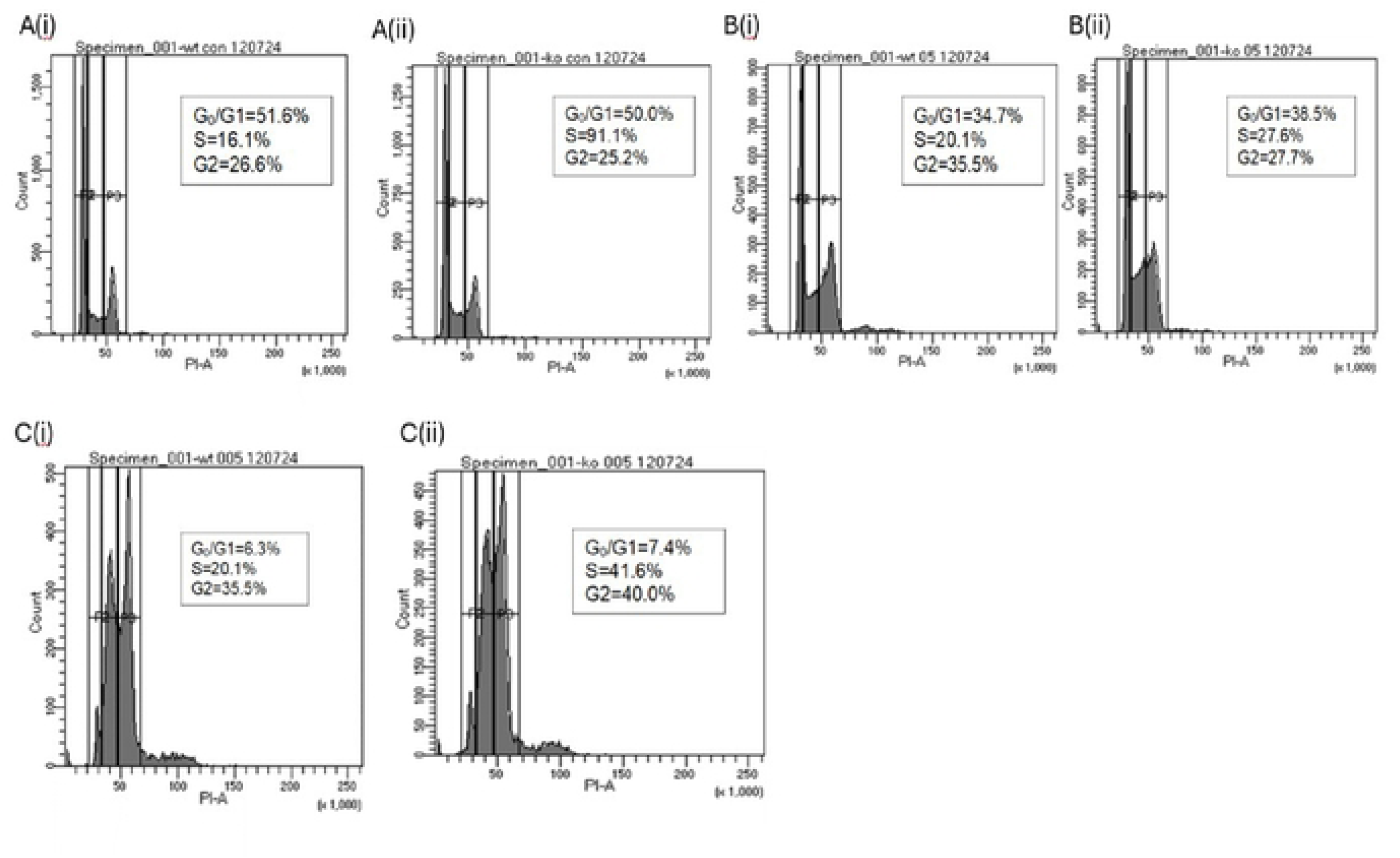
Cell-cycle distribution of WT and ZFP36L1 knockout MDA-MB-231 cells following exposure to different concentrations of doxorubicin. Representative flow cytometric analysis of WT and ZFP36L1 knockout (KO) MDA-MB-231 cells following treatment with 0.5 μM and 0.005 μM doxorubicin for 24 h. Histograms illustrate the distribution of cells within the G₀/G₁, S, and G_2_/M phases following DNA staining with propidium iodide. The X-axis represents DNA content, while the Y-axis represents cell count. Panel A(i) shows WT cells before DOX treatment; (ii) shows KO cells before DOX treatment, B(i) shows WT cells after treatment with 0.5 μM DOX; Bii) shows KO cells after treatment with 0.5 μM DOX and C(i) shows WT cells after treatment with 0.005 μM DOX; (ii) shows KO cells after treatment with 0.005 μM DOX

### 3.4 Loss of ZFP36L1 is associated with increased DNA damage following doxorubicin treatment

DNA damage was evaluated using γ-H2AX flow cytometric analysis following treatment of WT and ZFP36L1 knockout (KO) MDA-MB-231 cells with 0.5 μM doxorubicin for 24 h. DOX treatment increased γ-H2AX fluorescence intensity in both WT and KO cells, indicating induction of DNA double-strand breaks (Figure 4).

**Figure 4.**
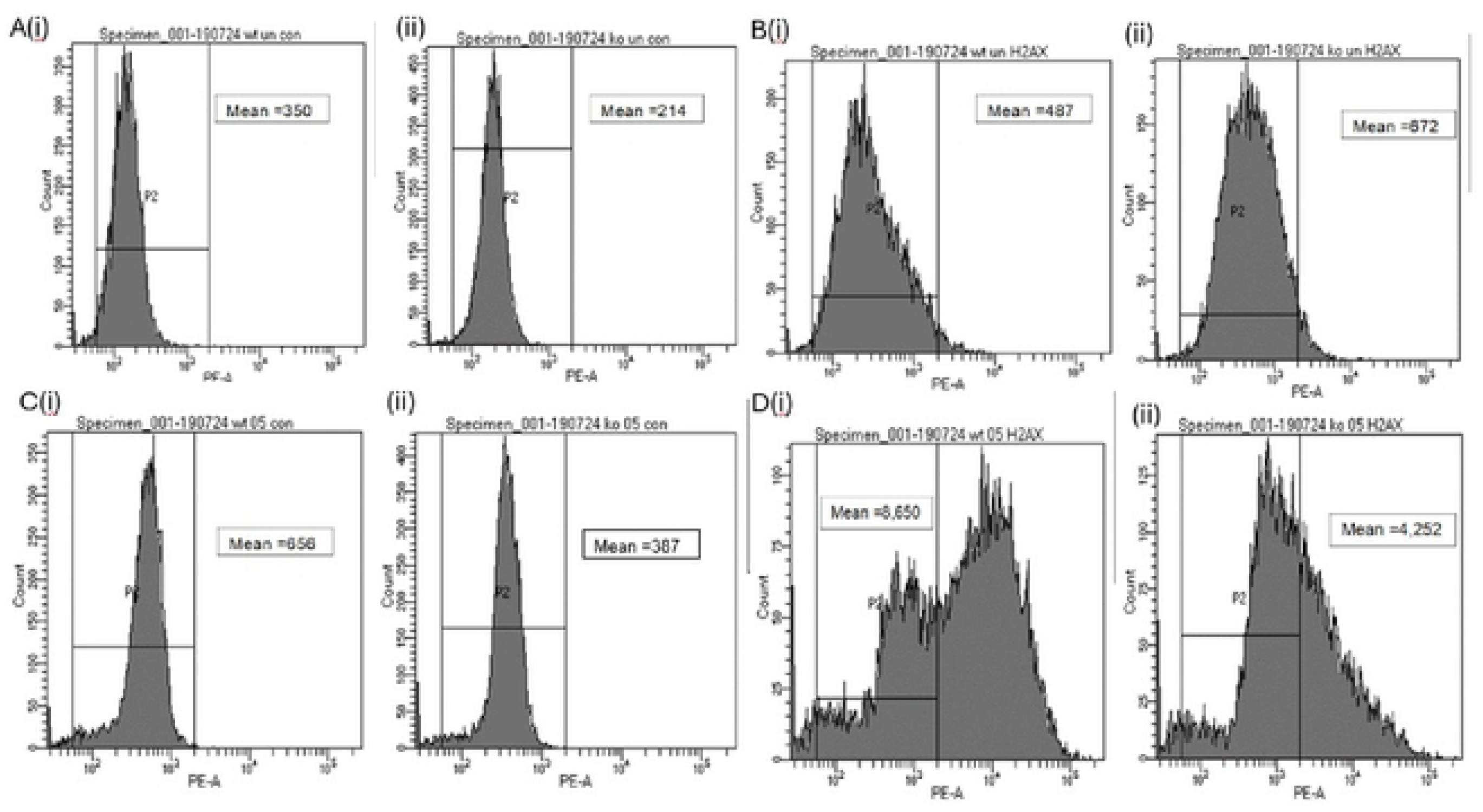
γ-H2AX expression in WT and ZFP36L1 knockout MDA-MB-231 cells following doxorubicin treatment. Representative flow cytometric analysis of γ-H2AX expression in WT and ZFP36L1 knockout (KO) MDA-MB-231 cells before and after treatment with 0.5 μM doxorubicin for 24 h. Cells were stained with anti-γ-H2AX antibody and analyzed in the PE fluorescence channel. Histograms represent γ-H2AX fluorescence intensity (x-axis) and cell count (y-axis). Control antibody-stained samples were included to establish background fluorescence. Panel A(i) shows WT cells untreated and stained with the control antibody, and A(ii) shows KO cells untreated and stained with the control antibody. Panel B(i) presents WT cells untreated and stained with the γ-H2AX antibody, while B(ii) shows KO cells untreated and stained with the γ-H2AX antibody. Panel C(i) displays WT cells treated with 0.5 µM DOX and stained with the control antibody, and C(ii) shows KO cells treated with 0.5 µM DOX and stained with the control antibody. Panel D(i) depicts WT cells treated with 0.5 µM DOX and stained with the γ-H2AX antibody, while D(ii) presents KO cells treated with 0.5 µM DOX and stained with the γ-H2AX antibody.

Compared with WT cells, untreated KO cells exhibited a rightward shift in γ-H2AX fluorescence intensity together with a higher mean fluorescence intensity, indicating elevated basal levels of DNA damage. Following DOX exposure, γ-H2AX expression increased further in both cell lines; however, the increase remained more pronounced in KO cells than in WT cells (Figure 4). These findings demonstrate that ZFP36L1-deficient cells exhibit increased basal DNA damage and an enhanced DNA damage response following exposure to DOX.

## 4. DISCUSSION

This study employed CRISPR-Cas9 mediated ZFP36L1 deletion in MDA-MB-231 TNBC cells to examine the functional significance of ZFP36L1 in breast cancer proliferation, doxorubicin chemosensitivity, cell cycle checkpoint regulation, and DNA damage maintenance. Our results indicate that the loss of ZFP36L1 markedly diminished baseline proliferative capacity, suggested a decrease in DOX sensitivity, modified cell cycle checkpoint responses under genotoxic stress, and increased constitutive DNA damage markers even without exogenous insult.

At first glance, the finding that ZFP36L1 KO cells grew more slowly and in lower numbers than wild-type cells over the course of three weeks seems to be in opposition to the current tumor suppressor paradigm, where loss of ZFP36L1 should theoretically increase proliferation via increased stabilization of oncogenic transcripts (Loh et al., 2020). However, this discovery is consistent with accumulating evidence for context-dependent and even conflicting ZFP36L1 function. ZFP36L1 knockout by CRISPR-Cas9 in chronic myeloid leukemia also led to decreased cell expansion and increased expression of the cyclin-dependent kinase inhibitor CDKN1A via direct binding to the 3′UTR, leading the authors to state that ZFP36L1 cannot be definitively classified as a tumor suppressor but rather has an ambiguous, cell-type-specific role (Kaehler et al., 2021). The lower proliferation of ZFP36L1 KO cells in the current TNBC model could also be due to the compensatory overexpression of cell cycle inhibitors or checkpoint stalling caused by persistent replication stress, as demonstrated by the highly raised baseline γH2AX signal in untreated KO cells.

This ambiguity is further evidenced by recent work in muscle-invasive bladder cancer, where ZFP36L1 reduces colony formation and self-renewal, but also promotes epithelial-mesenchymal transition and invasion via metastasis-associated pathways (Yuan et al., 2022). Overexpression of ZFP36L1 in osteosarcoma also impaired non-homologous end joining (NHEJ) DNA repair by destabilizing DCLRE1C mRNA, hence increasing chemotherapy-induced DNA damage and death (Zhuang et al., 2025). These contradicting results emphasize that the tumor suppressive or oncogenic activity of ZFP36L1 is dependent on the cellular context and signaling environment and necessitates functional characterization in a cancer type-specific manner.

Furthermore, we found a high basal level of γH2AX in untreated ZFP36L1 KO cells. Phosphorylation of γH2AX at serine 139 is classically linked with DNA double strand breaks, but is becoming appreciated as a sensitive sign of replication stress that may occur prior to overt DNA breaking and can be mediated by activation of ATR-dependent kinases at stalled forks (Saini et al., 2020). In the absence of exogenous DNA damage, ZFP36L1 KO cells displayed a right-shifted γH2AX peak and higher mean fluorescence intensity than wild-type cells, suggesting that ZFP36L1 is necessary for the maintenance of replication fork stability and genomic integrity in rapidly proliferating breast cancer cells. This is consistent with recent evidence that ZFP36L1 overexpression inhibits non-homologous end joining (NHEJ) DNA repair by destabilizing DCLRE1C mRNA, hence aggravating chemotherapy-induced DNA damage and death in osteosarcoma (Zhuang et al., 2025). Alternatively, the lack of ZFP36L1 might interfere with the appropriate coordination of basal DNA repair, resulting in a buildup of unresolved replication intermediates, persistent low-level DNA damage signaling, and a resulting proliferative delay.

The ZFP36L1 KO cells had a tendency to greater IC50s (lower sensitivity) to doxorubicin than the wild-type cells, although the difference was not statistically significant (p=0.569). Thus, ZFP36L1 may regulate DNA damage response, but is not the major predictor of DOX sensitivity in the MDA-MB-231 context. The lack of statistical significance may be due to the absence of statistical power for the sample size (n=3–4), variability between experiments or functional redundancy with other members of the TTP family such as ZFP36L2 or TTP itself which may compensate for ZFP36L1 loss in regulating ARE-containing transcripts involved in drug resistance (Saini et al., 2020). Recent research supports the redundant idea. In osteosarcoma, ZFP36 and ZFP36L2 could both increase methotrexate sensitivity, whereas co-overexpression of ZFP36L1 and ZFP36L2 could somewhat improve cell sensitivity, indicating possible compensation in the ZFP36 family (Zhuang et al., 2025). Furthermore, the simultaneous ablation of ZFP36L1 and ZFP36L2 in murine lymphocytes results in T-cell acute lymphoblastic leukemia through Notch1 stabilization, demonstrating the critical and largely overlapping functions of these paralogs in maintaining hematological genomic homeostasis (Saini et al., 2020). Functional redundancy may buffer phenotypic effects of ZFP36L1 single KO when combined with TNBC, particularly in acute drug sensitivity testing.

Interestingly, at lower concentrations (0.14–1.13 µM) DOX paradoxically boosted cell viability relative to untreated controls in both genotypes, which has been previously documented in TNBC and attributed to hermetic stress responses or activation of pro-survival autophagy at sub-lethal levels. The more pronounced loss of viability observed in wild-type cells at higher DOX concentrations, opens up the possibility that ZFP36L1 expression may sensitize cells to acute genotoxic stress, presumably by pushing the balance between DNA repair and apoptotic commitment.

Cell cycle analysis showed that DOX administration caused a dose-dependent redistribution of G0/G1 cells into S and G2/M phases in both wild-type and KO cells, consistent with DNA damage-induced checkpoint arrest. Notably, KO cells showed a more potent S and G2/M arrest at low DOX concentrations (0.005 µM), suggesting that deletion of ZFP36L1 may affect checkpoint resolution or the kinetics of cell cycle re-entry after genotoxic treatment. This is consistent with the known function of ZFP36L1 as a regulator of cell cycle transcripts such as CCND1 and E2F1 on the post-transcriptional level (Loh et al., 2020) and its unclear regulation of CDKN1A (Kaehler et al., 2021). The enhanced arrest phenotype in KO cells may represent a compensatory mechanism to overcome the accumulated replication stress in the face of genotoxic pressure or alternatively, a deficient checkpoint adaptation caused by altered degradation of cell cycle regulating mRNAs. Recently, a genome-wide CRISPR-Cas9 screen in MDA-MB-231 cells showed that DNA damage and cell cycle regulation pathways were the most enriched pathways for genes that desensitize cells to doxorubicin, with CDC25B and NBN identified as common essential genes between cisplatin and doxorubicin resistance (Shao et al., 2025). These data jointly substantiate the concept that ZFP36L1 is a cell cycle and DNA damage response regulator operating in a wider gene network of checkpoint control genes that collectively define chemotherapeutic responses in TNBC.

## CONCLUSION

This study demonstrates that ZFP36L1 is necessary for optimal proliferative capacity in MDA-MB-231 breast cancer cells and that its loss induces constitutive replication stress and alters cell cycle checkpoint dynamics under doxorubicin-induced genotoxic stress. These findings support a complex, context-dependent role for ZFP36L1 in TNBC biology that extends beyond a simple tumor suppressor framework. Further investigation into the post-transcriptional networks regulated by ZFP36L1 in breast cancer will be essential for determining its potential as a prognostic biomarker or therapeutic target for modulating chemotherapy response.

